# Flickering white light stimulation at 60 Hz induces strong, widespread neural entrainment and synchrony in healthy subjects

**DOI:** 10.1101/2025.01.27.634699

**Authors:** MohammadAmin Alamalhoda, Friederike Leesch, Francesca Giovanetti, Eoghan Dunne, Giuseppina Pilloni, Mark Caffrey, Jack O’Keeffe, Alessandro Venturino, Maria Teresa Ferretti

## Abstract

**Background:** While the effects of 40 Hz externally-induced neural entrainment have been extensively described, little is known about 60 Hz entrainment in humans. Given the role of 60 Hz in cognition, neuroplasticity and neuropsychiatric disorders, this warrants further investigation.

**Objectives:** This pilot study characterizes, for the first time, the neural and somatic response to 60 Hz light entrainment in healthy volunteers, over a 3 week-period.

**Methods:** Fourteen volunteers were randomized to receive either 60 Hz flickering white light or constant light as sham (30-min sessions, for 3 weeks, 5 days a week). Neural entrainment was assessed with EEG on days 1, 5 and 19. Salivary cortisol and C-reactive protein (CRP) levels, measured with ELISA, assessed the somatic response to stimulation. Side effects and well-being were monitored via questionnaires.

**Results:** 60 Hz flickering light induced a strong neural entrainment across visual, parietal, temporal and frontal cortex. The signal was highly synchronous but declined significantly by day 19 compared to day 1, indicating neural habituation. Cortisol and CRP salivary levels were unchanged and the stimulation was well tolerated.

**Conclusions:** To the best of our knowledge this is the first study to characterize both the neural and the somatic response to flickering light over 3 weeks. The observed neural habituation suggests that neuroplasticity could be induced with repeated stimulations over 3 weeks. 60 Hz stimulation for modulating brain activity and induce neuroplasticity has implications for our basic understanding of brain physiology as well as treatment of psychiatric disorders.

## 1. Introduction

*Neuromodulation* is a collection of techniques aimed at modulating diffuse neuronal activity to achieve therapeutic effects. When modulation is obtained through the application of external energy (e.g., electrical currents, magnetic field, light, or ultrasound) to the brain, it is referred to as *neurostimulation*^1^. Non-invasive brain stimulation (NIBSs) techniques, such as electroconvulsive therapy (ECT), transcranial magnetic stimulation (TMS), and transcranial electrical stimulation (tES), exert measurable structural and functional effects on the brain. These effects include increased neuroplasticity^2^, changes in brain structure^3^ and connectivity^4^, and restoration of brain-derived neurotrophic factor (BDNF) levels^5^. Some NIBSs solutions are FDA-approved for the treatment of various brain diseases, including depression^6^ and obsessive-compulsive disorder^7^ (OCD), underscoring the therapeutic potential of neuromodulation.

*Emerging non-invasive neuromodulatory solutions* leverage different energy modalities to modify neuronal excitability and modulate brain activity. These include ultrasounds, such as transcranial focused ultrasound^8^; interference of electric fields, such as temporal interference stimulation^9^; and thermal stimulation using near-infrared lasers^10^.

*A new promising, fully non-invasive brain stimulation technique utilizes intermittent sensory stimulation*. Brainwaves naturally synchronize with the rhythm of periodic external stimuli, such as flickering lights, speech, music, or tactile stimuli^11^. The synchronization of brainwaves is known as entrainment. Synchronization in the gamma range (between 30 and 70 Hz) is of particular therapeutic interest, as gamma brainwaves occur naturally when the brain is concentrated and involved in cognitive and executive functions^12^, and their alterations have been described in several neuropsychiatric conditions^13,14^.

*Multisensory (combined visual and acoustic) external stimulation* at 40 Hz has been extensively studied in both mouse models^15^ and humans^16^. This stimulation has been shown to induce strong and well-tolerated brain entrainment. Initial findings suggest its potential for treating Alzheimer’s disease^17^; in particular, 40 Hz stimulation increased connectivity between the precuneus and posterior cingulate cortex in Alzheimer’s patients, indicating neuroplastic effects mediated by brain entrainment^18^.

In contrast, the effects of 60Hz remain largely unexplored in humans, despite evidence that this frequency plays a role in cognitive and executive functions and is altered in neurological and psychiatric conditions^13,14^, this area warrants further investigation. Our previous preclinical work in mice demonstrated that visual 60 Hz stimulation with flickering light in mice induces neural entrainment and juvenile-like neuroplasticity through microglia-mediated remodeling of the perineuronal nets (PNNs)^19^. However, 60 Hz light entrainment in humans has not been extensively studied, particularly regarding the effect of repeated daily stimulations over time.

*The scope of this pilot study* is to characterize the effect of 60 Hz light-based stimulation on healthy human brains; we believe this is the first study to provide such characterization from a single session to a duration of 3 weeks. We applied LED-generated, 60 Hz pulsed flickering white light with a square wave modulation to a group of 14 healthy volunteers. The acute (single session), short-term (after 5 days), and intermediate (after 3 weeks) effects on brain electrical activity were assessed from electroencephalogram (EEG) data and compared to the sham stimulation of constant light. Salivary levels of biochemical markers –cortisol and C-reactive protein (CRP)– and participant questionnaires were used to monitor the somatic responses and tolerability. Our findings show that 60 Hz flickering light stimulation induces strong and widespread neural entrainment, with a high level of synchronization across brain regions. Notably, the entrainment and synchronization adapt over time, suggesting a neuroplastic response induced by 60 Hz. The stimulation is well tolerated by healthy individuals.

## 2. Methods

In this section the recruitment of participants, the experimental design, the device setup and the analysis performed will be presented.

### 2.1 Participants

The study involved a cohort of 14 young adult healthy volunteers (demographics in **Table 1**). The primary exclusion criteria were diagnosis of neurodegenerative or psychiatric diseases, in particular history of seizures or epilepsy, migraine, and tinnitus. Full inclusion and exclusion criteria are listed in the **Supplementary Table 1**.

**Table 1.** Demographic of study participants. The table summarizes the main characteristics of the study participants. Sex was self-reported. Data were analyzed using a t-test for years of education, a Wilcoxon U test for age and Fisher’s exact test for proportions. No statistically significant differences were observed across groups.

| Characteristic | Sham group<br>(n=6) | Active group<br>(n= 8) | p-value |
| --- | --- | --- | --- |
| Age in years, mean ± SD | 25.5 ± 2.59 | 28.63 ± 6.37 | 0.6 |
| Sex, female no (%) | 3 (50) | 5 (62.5) | 1 |
| Education in years, mean ± SD | 19.63 ± 3.78 | 18.83 ± 3.19 | 0.68 |
| Race, no (%) |  |  |  |
| Caucasian | 4 (66.67) | 8 (100) | 0.16 |
| Asian | 2 (33.33) | 0 (0) |  |

The 14 participants were randomized to active stimulation (n=8) or sham (n=6). The groups were balanced in terms of self-reported sex, age, and education (**Table 1**). Two participants in the active group dropped out for personal reasons (one completed one week of stimulation, and the other completed 16 days). The EEG data of these two individuals who dropped out were not included in the analysis because of missing data; similarly, these two samples were excluded from the biochemical analysis because of high viscosity. Both individuals completed the final questionnaire and therefore their questionnaire and side effects data were analysed.

### 2.2 Ethical Approval and Overall Study Design

The study was conducted in accordance with the World Medical Association Helsinki Declaration for human experimentation. All procedures were performed in compliance with relevant laws and have been approved by the Lower Austria Ethical Commission (study GS3-EK-4/908-2024 approved on July 1^st^ 2024). All procedures were carried out at Plöcking 1, XISTA Science Park, 3400 Klosterneuburg, Austria, under medical supervision. Written consent was obtained by study personnel from all participants. After the assessment of eligibility, participants were randomly assigned with equal probability (0.5) to either the active stimulation group (60 Hz white light) or the sham group (constant light; **Fig 1B**). The study was single-blind, where participants were not aware of their group assignment. To check the effectiveness of the blindness procedure, at the end of the study, participants were asked to guess which stimulation they received and on average 60% of them, in both groups, guessed correctly (**Table 2**).

**Figure 1.**
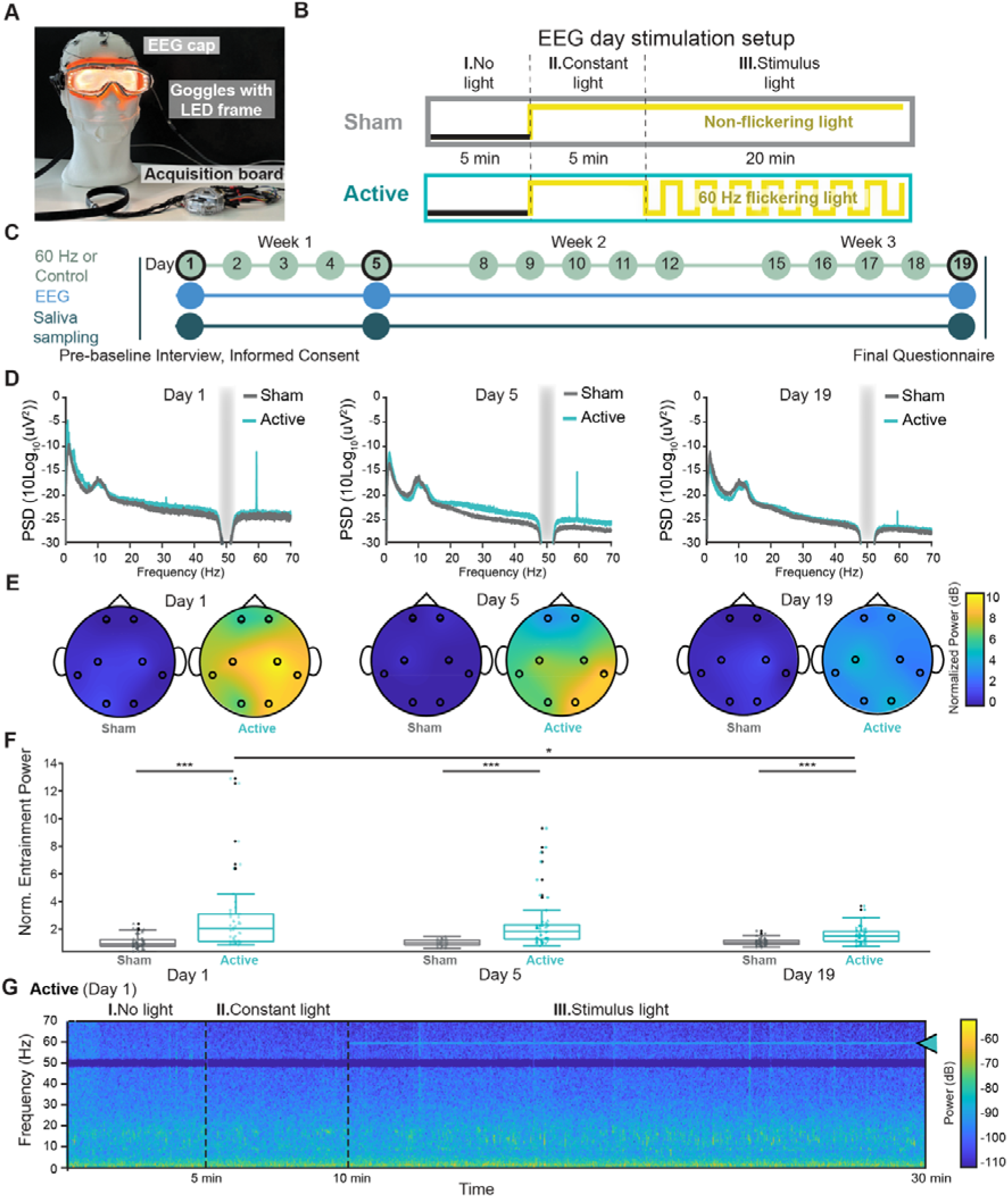
60 Hz-induced neural entrainment. **(A**) Schematic of the experimental setup, showing a wearable device (adapted from a Uvex mask equipped with LED lights) designed to be compatible with an EEG setup. **(B**) Overview of each EEG session, which included two control phases (I. no light and II. constant light) and one stimulus phase (III. stimulus light (**C**) Diagram of the experimental timeline: participants underwent light stimulation over three weeks, with EEG recordings and saliva sampling performed on days 1, 5, and 19 (indicated by black circles). On EEG days, the stimulation setup depicted in Panel B was applied. On non-EEG days (no black circles), participants received either an active stimulus (60 Hz flickering light) or a sham stimulus (constant light) for 30 minutes per session. **(D**) Scalp EEG power spectral density (PSD) averaged across all channels for participants in each group. The gray bar indicates the 50 Hz line noise, which was notch-filtered. A strong 60 Hz frequency component is evident from start to finish, but decreases over the course of the study at day 5 and day 19. (**E**) Topographic maps showing normalized changes in 60 Hz PSD (relative to baseline) averaged across participants of each group. (**F**) Significant differences in normalized changes in 60 Hz PSD values were observed across all channels between the active and sham groups on days 1, 5, and 19. Statistical significance for these inter-group comparisons was assessed using the Wilcoxon rank-sum test. Furthermore, within the active group, significant differences in normalized PSD were detected between day 1 vs. day 5 and day 1 vs. day 19, accounting for repeated measurements. These intra-group comparisons across days were evaluated using the Kruskal-Wallis test, followed by post-hoc pairwise comparisons performed using Dunn’s test with Bonferroni correction. The normality of the data was assessed using the Shapiro-Wilk test, and non-parametric methods were applied due to deviations from normality. In this figure, different channels are shape-coded. * = p < 0.05, *** = p < 0.001. Detailed p-values are provided in **Supplementary Table 2**. **(G**) Short-Time Fourier Transform (STFT) of a representative active group participant, demonstrating visible 60 Hz entrainment during light stimulation, as indicated by the arrow.

**Table 2.**
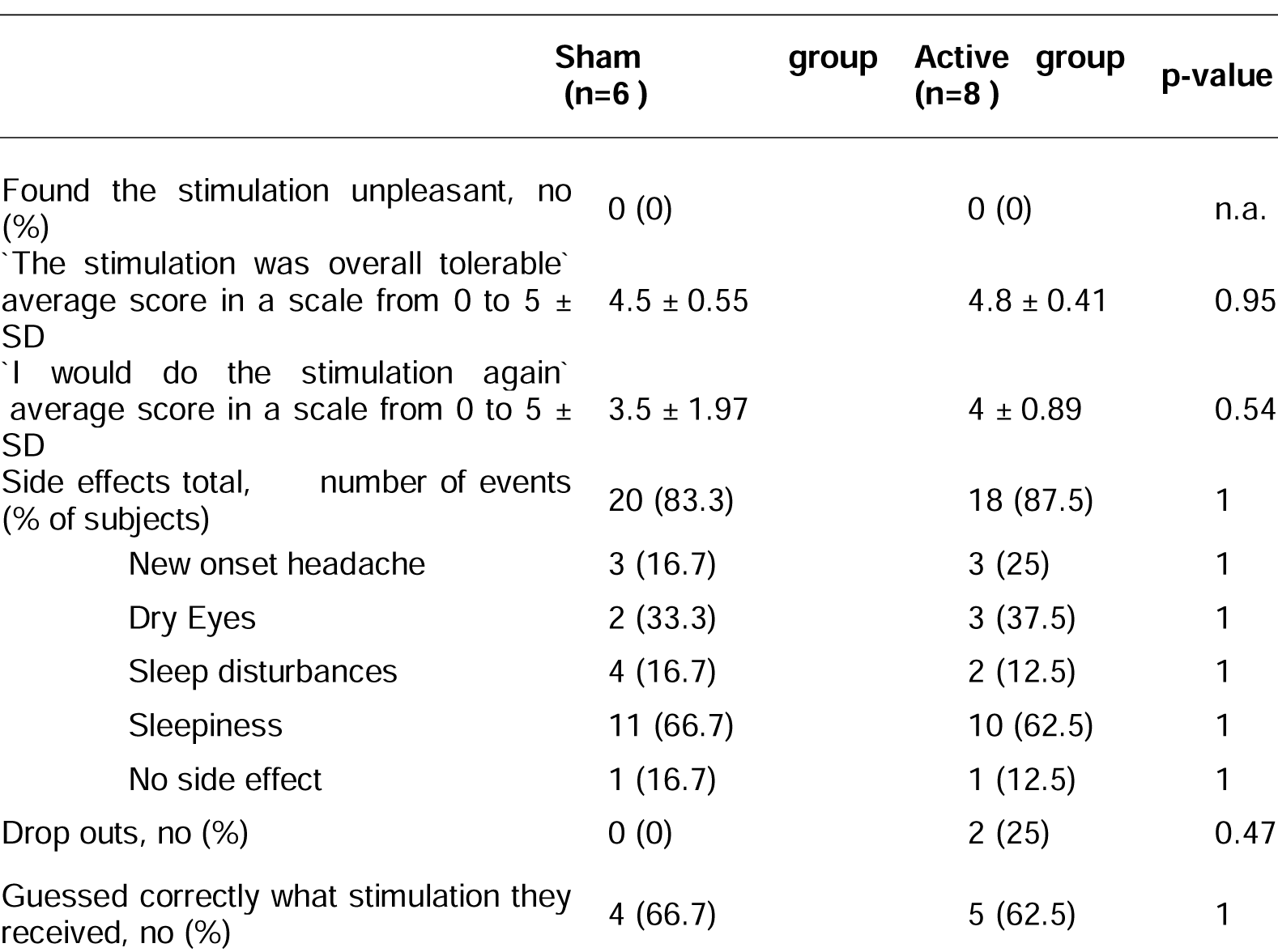
Safety and Tolerability. The Table shows the analysis of data collected from questionnaires (at the end of the study) and self-reports (throughout the study) to assess tolerability and safety of 60 Hz stimulation (active) as compared to stimulation with constant light (sham). Participants rated statements on a scale from 0 (‘ not at all’) to 5 (‘absolutely’). Both light stimulation conditions, sham and active, were well tolerated and induced only minor side effects. Discrete data were analyzed using the Mann-Whintney-Wilcoxon U test, while all other data were analysed with Fisher’s exact test. No statistically significant differences were observed between the sham and active groups.

Each participant received the stimulation 5 days a week, Monday to Friday, for 3 consecutive weeks, according to the light stimulation protocol described below (**Fig 1C**). The stimulation sessions took place during light-time, between 9 am and 4 pm. On days 1, 5, and 19 an EEG recording was performed during the stimulation and a saliva sample was collected. The time of delivering the stimulation for each subject remained consistent across all days. Side effects were collected daily based on self-report; a final questionnaire was administered at the end of the study to investigate tolerability, side effects, and blinding. All data were pseudonymized and handled according to national and international laws; the privacy rights of human subjects have been observed.

### 2.3 Light Stimulation Device Design

To deliver flickering white light stimulation in a controllable and standardized manner, we developed a wearable headset derived from Uvex Ultrasonic safety glasses (Manufacturer Part No 9302245). The device was worn around the face and head, resembling safety glasses. A strip of LEDs (PowerLED Chromatic LED strip - Manufacturer Part No D0-55-35-1-120-F8-20-FP) was integrated around the ocular component to direct light tangentially to the periphery of the wearer’s eyes (see **Fig 1A**). This LED strip has 120 LEDs per meter, and 35 cm of this LED strip was used in the headset.

The LEDs emitted a square-wave function of white light at 60 Hz with a duty cycle of 50% (rise time: 1.5 ms, fall time: 3.5 ms). The stimulation frequency was manually measured using an oscilloscope (Keysight Edux1052A, Manufacturer Part No 302-25-092) before each data collection to ensure consistent frequency throughout the recording. The target frequency was set at 59.6 Hz (a 0.6% deviation from 60 Hz) to avoid the challenges associated with generating an exact 60 Hz signal. Maintaining 60 Hz required our microcontroller to toggle every 16.666… milliseconds, which could not be precisely represented due to the repeating decimal. This limitation could result in cumulative rounding errors and frequency drift over time. By using 59.6 Hz, the period (16.778 ms) could be represented more accurately, ensuring consistent stimulation frequency over the course of the stimulation. The light spectrum ranged from 440□nm to 770□nm (with full spectrum of visible light colours), corresponding to daylight wavelengths, measured with a spectrometer (ThorLabs CCS200; **Supplementary fig 1A**).

The color temperature was matched at 4000°K, and the light intensity was adjustable between 40 and 110 μW (measured at 2cm distance from the LED strip using ThorLabs PM100D); this range of intensities is considered safe to the eyes. The exact intensity was selected individually per participant to ensure a comfortable experience during each session based on the participant’s verbal feedback. On average, subjects chose a light intensity of 80 μW (STD: 7 μW across subjects), and this intensity remained largely consistent over the days of stimulation for each subject (maximum STD: 9 μW for any individual subject).

To minimize electrical noise interference with the EEG setup, the LED strip was enclosed within a copper mesh (Thorlabs, Catalog No. PSY406), and powered using a long shielded cable (RS PRO data cable - Manufacturer Part No 303-95-459), both grounded to the EEG acquisition board, effectively creating a Faraday cage to eliminate electromagnetic emissions from both the powering cable and the LED strip. The efficacy of the shielding was measured at a distance of 10 cm from the LED strip using an EMF meter (MULTIFIELD-EMF450). The magnetic field measured with LED on was 203 uT without the cage, and it reduced to 66 uT when using the cage.

The 60 Hz band square signal was generated using an Arduino Nano board PWM pin (Arduino, Catalog No. SKU A000005) soldered onto a Printed Circuit Board (Manufacturer Part No RE942-S3), connected to the LEDs via a circuit that included a 10 kΩ potentiometer (Manufacturer Part No P16NP103MAB15) for intensity modulation and a p-channel MOSFET (Manufacturer Part No IRF9540NPBF) to drive sufficient current to the LEDs. The gate of the MOSFET was driven using a MOSFET gate driver (Manufacturer Part No TC4420CPA), to ensure fast switching of the MOSFET transistor. The entire setup was powered using a power bank (Manufacturer Part No 57976 101 111) placed more than 1 m away from the subject. Additionally, the prototype was equipped with a photoresistor (Manufacturer Part No TS2134-A) to capture the light emitted from the LEDs for synchronization of the onset of the light stimulation with the EEG signals during data processing. The photoresistor was connected to an Arduino Nano board, where its analog signal was converted into a digital signal. This digital signal was then transmitted to one of the digital pins on the acquisition board.

### 2.4 Light Stimulation Protocol (active and sham)

Participants were seated comfortably on a chair with a headrest to provide additional support and ensure stability. The wearable device was carefully positioned on the participant’s face to ensure both comfort and secure placement. The environment was kept quiet, with dim ambient lighting (**Supplementary fig 1B**). The operator activated the controller and adjusted the light intensity based on real-time verbal feedback from the participant to maximize comfort. The light adjustment was performed during the first minute of the stimulation. This first minute was then excluded from the EEG signal. For active stimulation, on each of the EEG recording days (day 1, day 5, and day 19), the experimental protocol consisted of three distinct periods (**Fig 1B**):

1. Baseline (No light): A 5-minute initial period without light exposure.
2. Constant Light Exposure (Constant light): A subsequent 5-minute period of constant light exposure.
3. Stimulus-Modulated Light Exposure (Stimulus light): A 20-minute final period of stimulus-modulated light exposure (60 Hz)

On all non EEG-days, the protocol was simplified to a continuous 30-minute session of 60 Hz light exposure.

For the sham stimulation, the exact same procedure and device were used. On EEG recording days (day 1, 5 and 19) volunteers underwent the 3-step protocol described above but for step III they received constant light as stimulus light. On all other days, they received 30 minutes of constant light.

Each participant (active and sham) received the stimulation daily, Monday through Friday, for three consecutive weeks, for a total of 15 sessions each. Once started, all stimulation sessions (active and sham) were completed and there were no cases of volunteers asking to interrupt beforehand.

### 2.5 EEG Recording and data capture

EEG data were recorded using the Cyton + Daisy Biosensing Boards (OpenBCI; 10 electrodes at 250Hz sampling rate - 8 active electrodes and 2 reference electrodes) - PIC32MX250F128B microcontroller) and the OpenBCI EEG Cap Kit (OpenBCI, 19-channel cap), medium size, fitted with sintered wet electrodes (sintered Ag/AgCl coated electrodes). Sintered wet electrodes were selected due to their known low impedance and stable signal acquisition properties, ensuring high-quality data capture over repeated use. Ten electrodes were placed according to the internationally recognized 10–20 system^20^, ensuring consistent electrode positioning for comparability across studies. In this study, we specifically used channels FP1, FP2, C3, C5, T5, T6, O1, and O2 targeting key frontal, parietal, temporal, and occipital regions (**Supplementary fig 1C**). A low-impedance electrode gel (OpenBCI 380130) was applied using a syringe with a blunt needle to ensure very low-impedance electrical contact between the scalp and the electrodes. The impedance of all channels were kept under 5 kΩ during all the recordings. The EEG signals were referenced to the right earlobe to establish a stable baseline for differential measurements. To minimize environmental noise and enhance the signal-to-noise ratio, the EEG system’s bias electrode was connected to the subject’s left wrist. This configuration allowed the differential amplifier to effectively reject common-mode noise by accounting for the bias voltage (*V_b_*) in signal processing. The amplified EEG signal was obtained using the equation I:

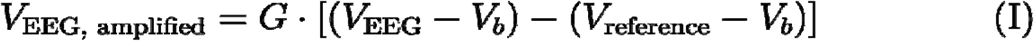

where, *V_EEG_* represents the raw EEG signal, *V_reference_* is the reference signal from the right earlobe, *V_b_* is the bias voltage, and G denotes the gain of the amplifier. By integrating the bias voltage into both signal paths, the system effectively eliminated common-mode interference, yielding a cleaner and more accurate representation of the neural activity for subsequent analysis.

Data was wirelessly transmitted from the acquisition boards to a computer via the Cyton board’s built-in radio module (over Bluetooth), minimizing cable artifacts and enhancing participant comfort during recording sessions. The board was powered by a rechargeable lithium battery (Manufacturer Part No 304-24-383 - 5 V, 2A output), providing a stable power supply throughout the experiment. The recordings were performed using the OpenBCI GUI software (version 6.0.0-beta.1) on an Apple MacBook Pro computer with an M2 processor and 16 GB of RAM. The EEG cap was cleared after each use using a brush to remove the residual gel off the cap’s electrodes, and was soaked in warm water for 15 minutes so that the remaining gel dissolves quickly. The cap was fully dried afterwards. To disinfect the cap, it was soaked for up to 30 minutes in a diluted bleach solution containing approximately 100 ppm sodium hypochlorite, then rinsed with clean water and hung to dry completely. The sintered Ag/AgCl electrodes are resistant to corrosion and the cap was manually inspected daily.

### 2.6 Signal Processing and Analysis

#### EEG Data Preprocessing

Since two subjects in the active group dropped out for personal reasons (one in week 1 and one in week 2), we decided to analyze only the data from the 6 subjects in this group who completed the study. EEG data preprocessing was performed using MATLAB (MathWorks Inc., Natick, MA) and the EEGLAB toolbox^21^. The raw signals were first notch-filtered at 50Hz to remove power line noise, followed by a bandpass filter from 0.5 Hz to 80 Hz using a butterworth 2nd order filter, applied bi-directionally to the signal to retain relevant EEG frequencies. The signals were then averaged and re-referenced to eliminate common noise across electrodes. Then, using the signal captured via the photoresistor, the signals were divided into baseline (5 minutes), constant light (5 minutes), and stimulus light (20 minutes) for further analysis. After, segments of data containing excessive noise or artifacts were manually inspected and removed. Finally, Independent Component Analysis (ICA) was applied to the preprocessed data to remove ocular and muscle movement artifacts^22^. Subsequent analyses were conducted using the MATLAB EEGLAB toolbox (MATLAB R2024b, and EEGLab v2024.2.1). Additional analysis and visualization were performed with open-source software ElecPhys (Github.com/AminAlam/ElecPhys). The custom code used for preprocessing data and analysis is available at Github.com/AminAlam/HVS.

#### EEG Data Analysis - Fast Fourier Transform (FFT)

The frequency content of the EEG signals was analyzed using the Fast Fourier Transform (FFT) implemented in MATLAB to obtain the PSD of the signals. The FFT is an efficient algorithm for computing the Discrete Fourier Transform (DFT) of a sequence, which transforms discrete time-domain signals into their frequency-domain representation^23^.

The DFT of a discrete signal *x*[*n*] of length N is defined as in equation II:

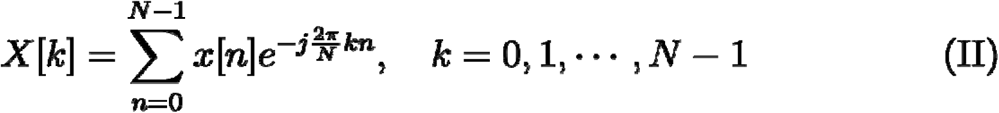

where *X*[*k*] is the complex amplitude of the *k^th^* frequency component, *x*[*n*] is the time-domain signal, *j* is the imaginary unit, and *N* is the number of samples. By applying the FFT, we were able to identify spectral components associated with the 60 Hz band light stimulation and assess changes in EEG power spectra across different experimental conditions.

#### EEG Data Analysis - Topoplots

For each light condition, the power spectral density (PSD) in the 60 Hz band (59.5 Hz to 60Hz) was normalized relative to the total power within the 2–80 Hz frequency range. Subsequently, the normalized PSD values for each light condition (constant light and stimulus light) were further normalized to the no-light condition to quantify the relative increase in 60 Hz band power compared to baseline. These final values were visualized as topoplots to illustrate the spatial distribution of the changes across the scalp.

#### EEG Data Analysis - Short-Time Fourier Transform (STFT)

Time-frequency analysis was conducted using the Short-Time Fourier Transform (STFT) with a Hamming window of 0.5-second length and 0.25-second overlap between windows. The STFT provides a way to analyze the frequency content of non-stationary signals over time by applying the Fourier Transform to short, overlapping segments of the signal^24^.

The STFT of a discrete-time signal *x*[*n*] is defined as in equation III:

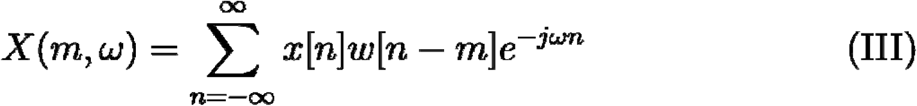

where *x*[*n*] is the input signal, *w*[*n*] is the window function (in this case, a Hamming window), *m* is the time index corresponding to the center of the window, *ω* is the angular frequency, *j* is the imaginary unit.

By sliding the window across the signal and computing the Fourier Transform at each position, we obtained a time-frequency representation that allowed us to observe transient changes in spectral power related to the light stimulation.

EEG Data Analysis - Phase-Locking Value (PLV). To assess the phase synchronization between EEG channels at the stimulation frequency, the Phase-Locking Value (PLV) was calculated. The EEG signals were filtered around 60 Hz (59.5 -60.5 Hz) using a 2nd order Butterworth filter applied bidirectionally, ensuring zero-phase distortion. The instantaneous phase of each signal was extracted using the Hilbert transform. The PLV between each pair of channels was computed using the formula25 shown in equation IV:

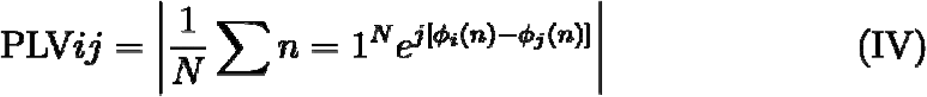

where *ϕ_i_*(*n*) and *ϕ_j_*(*n*) are the instantaneous phases of signals *i* and *j* at time point *n,N* is the number of time points, and *j* the imaginary unit.

The PLV for the all light conditions were normalized against the PLV of the no-light condition to quantify the relative increase in PLV compared to the baseline.

### 2.7 Biomarkers

#### Saliva collection

Saliva samples were obtained from participants on designated days (day 1, 5 and 19). Saliva samples were collected in 2 ml tubes (SARSTEDT, Cat. n. 72.695.500) following light stimulation, with the exact time of each collection carefully recorded. Participants were instructed to abstain from eating for a minimum of 1 hour before saliva collection and to avoid dairy products, caffeinated drinks, or acidic beverages within 15 minutes of sampling. Immediately following the collection, a Protease Inhibitor Cocktail (Sigma, Cat. n. #P2714) was added to the saliva at a concentration of 1 μL per 1 mL (v/v) of whole saliva, prepared according to the manufacturer’s instructions. Samples were then centrifuged at 2,000 g for 2 minutes to remove larger debris. The supernatant was aliquoted into 2 ml tubes, labeled with the participant ID code and date. Finally, samples were flash-frozen and stored at - 70°C for subsequent analysis.

#### Cortisol and CRP ELISAs and Analysis

Salivary cortisol and C-reactive protein (CRP) levels were measured using a competitive cortisol ELISA (Invitrogen, Cat. n. #EIAHCOR) and a quantitative CRP sandwich ELISA (Abcam, Cat. n. #ab108826). As a quality control step, samples with high viscosity, excessive debris, or mucus contamination were excluded from the analysis; high viscosity, which can be driven by dehydration, hormonal factors or bad dental status, was not compatible with pipetting26. To account for the strong diurnal variation of salivary cortisol levels, only samples collected within approximately the same time of the day (within 1h) were included in the cortisol analysis. Samples collected more than 1 hour apart were excluded due to reliability concerns. Ultimately, cortisol levels were analyzed in 9 participants (5 sham and 4 active), while CRP levels were analyzed in 10 participants (5 sham and 5 active).

The colorimetric reactions for both assays were measured using a Synergy H1 plate reader set to 450 nm, following the manufacturer’s protocols. CRP and cortisol concentrations were calculated using AssayFit Pro version 5.3.2 online analysis software. Relative fold changes in cortisol levels over the course of the study were determined by normalizing each time point to baseline (day 1) cortisol levels.

#### Safety, tolerability and blinding measures

To assess tolerability and gather general feedback on the stimulation, we designed a brief questionnaire and administered it to the participants at the end of the study. All participants (n=14) completed the questionnaire. The questionnaire contained 3 statements about the general experience:

1. ‘The stimulation was overall…’ -> pleasant/unpleasant
2. ‘The stimulation was overall tolerablè
3. ‘I would do the stimulation again’

For n.1, the participants were asked to select one option (pleasant/unpleasant). For statements n.2 and n.3 the participants rated them on a scale from 0 (not at all) to 5 (absolutely). The questionnaire also contained a question to check participantś blinding, by asking them: ‘Do you think you received the active or sham stimulation?’ (participants could select one option).

Adverse events were recorded daily based on spontaneous participants’ self-reports throughout the study.

### 2.9 Statistics

#### Statistical Analysis - Biomarkers and tolerability

Statistical comparisons between two groups were made using the two-sided T-test or Wilcoxon rank-sum U test due to the non-normal distribution of some data groups (tested using Shapiro-Wilk test). Wilcoxon rank-sum test was also used to analyze ordinal questionnaire data. Proportions were tested with Fisheŕs exact test. Significance was determined at p<0.05. Repeated measures over time were analyzed with two-way ANOVA. Statistical analyses were performed using GraphPad (versions 5 and 10.4).

#### Statistical Analysis - EEG entrainment power and PLV

Statistical analyses were performed using MATLAB to assess group differences and identify significant patterns in the data. Different statistical tests were applied depending on the nature of the data and the comparisons being made, with a focus on non-parametric methods due to violations of normality as determined by the Shapiro-Wilk test.

1) Inter-group comparisons (Control vs. Active groups):

To compare the Control and Active groups within each light condition and for each day, the Wilcoxon rank-sum test (also known as Mann-Whitney U test) was used. This test was chosen because it is a non-parametric method that does not assume normality and is appropriate for independent group comparisons. The rank-sum test evaluates whether the distributions of the two groups differ significantly. For each day within each light condition, the test provided p-values indicating whether there were significant differences between the Control and Active groups.

2) Intra-group comparisons across multiple days:

To evaluate differences across multiple days within each group (Control or Active) for a given light condition, the Kruskal-Wallis test was applied. This test is a non-parametric alternative to one-way ANOVA and is suitable for comparing more than two groups (in this case, days) when the assumption of normality is violated. The Kruskal-Wallis test assesses whether there are statistically significant differences in the distributions of the dependent variable across the days.

3) Post-hoc pairwise comparisons:

When the Kruskal-Wallis test indicated significant differences (p < 0.05), post-hoc pairwise comparisons were conducted using Dunn’s test with Bonferroni correction. This step was necessary to determine which specific pairs of days differed significantly. Dunn’s test adjusts for the increased risk of Type I errors when performing multiple comparisons, ensuring the results remain reliable.

## 4. Results

### 4.1 Participants and experimental setup used to study 60 Hz neural entrainment

Our primary goal was to assess whether 60 Hz light stimulation would elicit neural entrainment in healthy volunteers, as previously described in animal models^1^ and documented in humans at other frequencies, such as 40 Hz15.

To address this, we administered 60 Hz flickering light to a group of 8 volunteers (referred to as ‘activé group) and compared their response to a parallel group of 6 volunteers who received constant, non-flickering light (referred to as ‘sham’ group). No statistically significant differences were observed across groups in terms of sex, age, education and race, indicating that the random allocation generated two well-balanced groups (**Table 1**).

For the stimulation, we used a prototype wearable headset derived from safety glasses, lined with a strip of LEDs and equipped with a mini Faraday cage to minimize electrical noise (**Fig 1A, Supp.** Fig 1B). The stimulation lasted for 3 weeks, with participants undergoing one session per day from Monday to Friday (**Fig 1C**). Importantly, once a session was started, all participants completed it fully and we had no cases of interruptions or shortening of sessions. Two volunteers dropped out of the experiment before its completion for personal reasons; their EEG and biochemical data were not included in the EEG analysis below but their questionnaire and side effects data were analyzed.

Neural entrainment was investigated using an 8-channel EEG setup during the stimulation on days 1, 5 and 19. On EEG recording days, the stimulation followed a three-step experimental paradigm as illustrated in **Fig 1B**. Each subject sequentially underwent: (I.) an initial habituation phase of 5 minutes with no light, (II.) 5 minutes of constant light, and (III.) 20 minutes of stimulus light. For the sham group, the stimulus light (III.) consisted of non-flickering, constant light, while for the active group, it consisted of 60 Hz flickering light. The inclusion of the no-light condition and constant light condition served as critical controls to establish a robust baseline for interpreting the effects of stimulus light. The no-light condition was included to assess the participants’ baseline brain activity without any external visual stimulation, ensuring that any subsequent changes were attributable to light stimulation rather than inherent variability in brain activity. The constant light condition allowed for the isolation of general effects of light exposure independent of the flickering characteristics of the stimulus light. This approach provided a comprehensive framework to evaluate the specificity of neural entrainment to the 60 Hz flickering light.

### 4.2 60 Hz light induces neural entrainment

We investigated the occurrence and characteristics of neural entrainment at 60 Hz using different EEG signal analysis techniques. First, we performed a Fast Fourier Transform (FFT) analysis to investigate the frequency component of the EEG signal and calculate the power spectral density (PSD). If entrainment occurs, we expect to observe a peak at the administered frequency.

Indeed, the FFT analysis of EEG data from participants in the active group demonstrated a clear and distinct peak in the 60 Hz band during the stimulus-modulated light exposure on day 1 (**Fig 1D**). This peak was absent in the baseline period and in all conditions for the sham group (**Fig1D**, gray line, and **Supp.** Fig 1D).

The quantification of normalized entrainment power at 60 Hz band across individual channels (**Fig 1E, F**) confirms that the entrainment at 60 Hz band in the active group affected most channels in all subjects. The normalized power in the active group, averaged across channels per subject, was 2.8 on day 1 (SD 2.69), 2.4 on day 5 (SD 1.94) and 1.54 on day 19 (SD 0.61). The normalized power of individual channels ranged from 0.8 to 12.9 on day 1, from 0.7 to 9.3 on day 5 and from 0.7 to 3.6 on day 19.

On day 1 the normalized power in the active group across channels was significantly higher than in sham groups (**Fig 1F**). As shown in **Fig 1F** (with shape code per each subject) and **Supplementary** Fig 1H, while some variability was observed across subjects, most of them responded at day 1. Importantly, Time-frequency analysis using STFT revealed a consistent band of activity at 60 Hz band throughout the 20-minute stimulus in the active group (**Fig 1G**). This sustained 60 Hz activity was absent during the baseline period in the same participants and in the sham group (**Supplementary** Fig 1I). The continuous presence of the 60 Hz band over time strongly supports the effectiveness of the light stimulation in entraining neural oscillations at the desired frequency.

When analysing the effect over time, we observed a noticeable decrease in 60 Hz power on days 5 and 19 in the active group as compared to day 1, corresponding to a 14% and 45% decrease over day 1, respectively (**Fig 1D, E, F**). By day 19, the power in the active group was significantly reduced compared to day 1 in the same group (**Fig 1F**). However, the normalized power in the active group remained significant on each day as compared to the sham group (**Fig 1F**). Overall, these results indicate that 60 Hz light entrainment facilitates neural entrainment, with the oscillation observed across multiple brain regions and a gradual decrease in power over time.

### 4.3 The 60 Hz light entrainment is widespread and synchronized

Having confirmed the occurrence of neural entrainment, we next assessed its spatial distribution. Topographical maps of EEG power in the 60 Hz band (**Fig 1E, Supplementary** Fig. 1 **E**) showed a global increase in power across most electrodes on day 1 of stimulation in the active group. This increase was observed across frontal, parietal, temporal, and occipital regions, indicating widespread neural entrainment. Given the widespread pattern of brain activation, we performed phase-locking value (PLV) analysis to determine whether the 60 Hz neural entrainment observed across regions exhibited synchronization or consistent phase relationships. On day 1, strong phase synchronization was evident between most channel pairs in the active stimulation group (**Fig 2B, Supplementary fig 2B**), suggesting coherent neural activity induced by the light stimulation.

**Figure 2.**
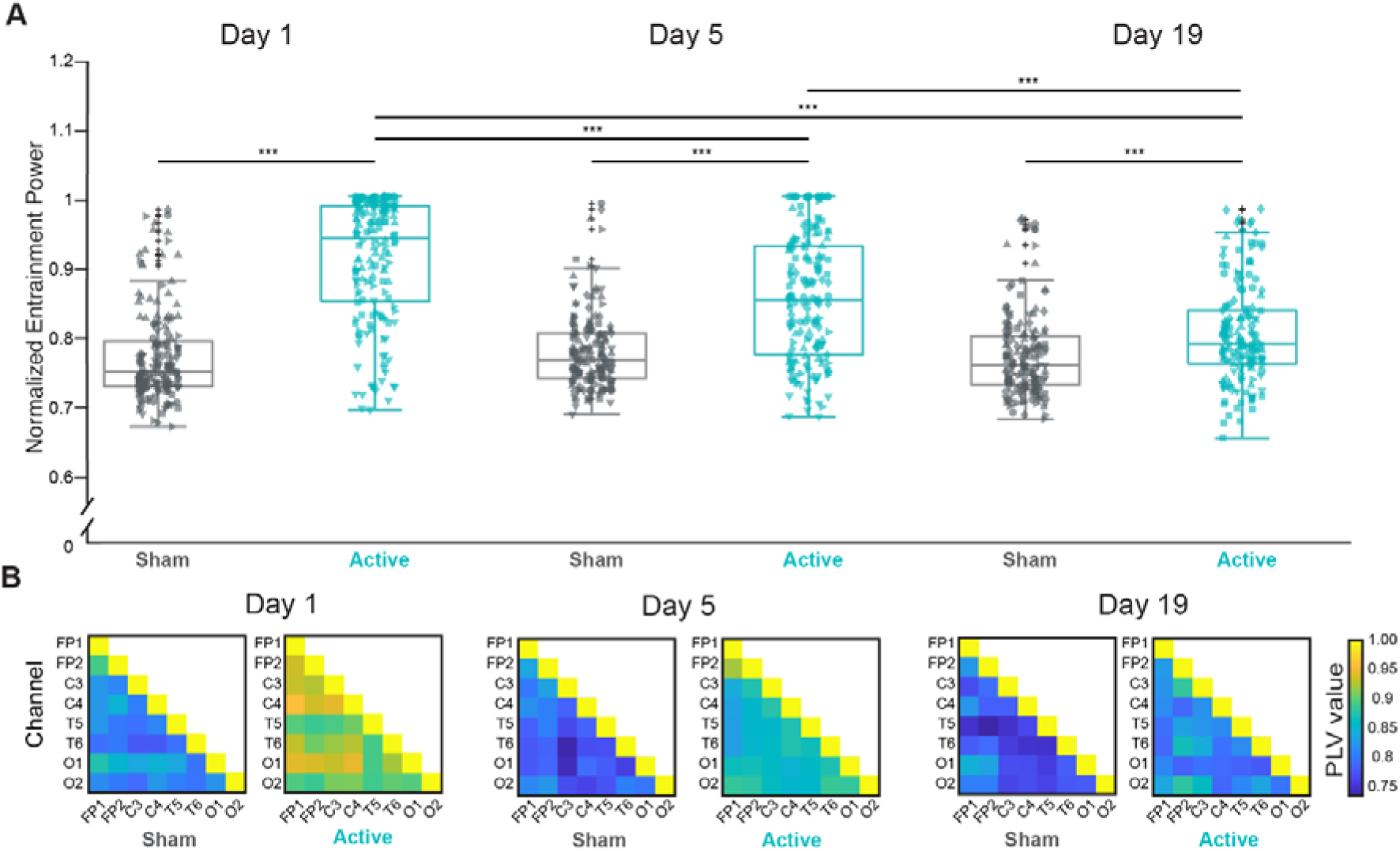
Synchronization of Brain Activity During 60 Hz Entrainment. Phase-Locking Value (PLV) analysis was performed to evaluate whether 60 Hz neural entrainment in the active group led to synchronization or consistent phase relationships across brain regions. Each PLV matrix was normalized to its corresponding baseline PLV matrix (PLV matrix during No Light) to highlight stimulation-induced changes. (**A**) Significant differences in normalized PLV were observed across all channel pairs between the active and sham groups on days 1, 5, and 19. Additionally, within the active group, significant differences in normalized PLV were detected between day 1 vs. day 5 and day 1 vs. day 19, accounting for repeated measurements. Statistical significance for these inter-group comparisons was assessed using the Wilcoxon rank-sum test. The intra-group comparisons across days were evaluated using the Kruskal-Wallis test, followed by post-hoc pairwise comparisons performed using Dunn’s test with Bonferroni correction. Normality of the data was assessed using the Shapiro-Wilk test, and non-parametric methods were employed due to deviations from normality. (**B**) PLV matrices for the active and sham groups across experimental days. Each element in the matrix represents the PLV for specific pairs of EEG channels, with diagonal elements showing a value of 1, indicating PLV between identical signals. * =p < 0.05, *** = p < 0.001. Detailed p-values are provided in **Supplementary Table 2.**

Statistical analysis confirmed robust and significant increase in PLV in the active group compared to the sham (**Fig 2A, Supplementary fig 2A**).

Similar to the changes observed in the power of entrainment, we detected a significant effect of time on the topographic distribution of power, with reductions observed on day 5 and day 19. Topographically, the power decline did not follow a consistent pattern. These results suggest a habituation effect or neural adaptation to repeated stimulations over time, as previously shown with other sensory modalities^27,28^.

Consistent with these findings, PLV significantly decreased from day 5 till day 19 (**Fig 2A**) in the active group. This reduction in PLV over time suggests a decline in neural synchrony at the stimulation frequency with repeated exposures.

### 4.4 Effects on cortisol levels measured in saliva

After having explored the neural response to 60 Hz flickering light, we wondered whether this stimulation might elicit somatic responses that can be measured with biomarkers. Various NIBS, including TMS^29^ and tDCS^30^, have been shown to lower cortisol levels in healthy participants, suggesting a potential effect of neuromodulation on the neuroendocrine axis. As far as we know this has not been investigated with light-induced neural entrainment, so we included this analysis in our experimental design. Salivary cortisol levels were presented as relative values, normalized to day 1 levels to account for sample variability (as done by^31^). Although no statistically significant differences were observed between sham and active participants in response to 60 Hz stimulation at day 5 or day 19 (**Fig. 3A**), active participants exhibited a notable trend toward reduced cortisol levels over the course of the stimulation. Future studies with a higher number of subjects might help elucidate whether 60 Hz neural entrainment has an effect on cortisol levels.

**Figure 3.**
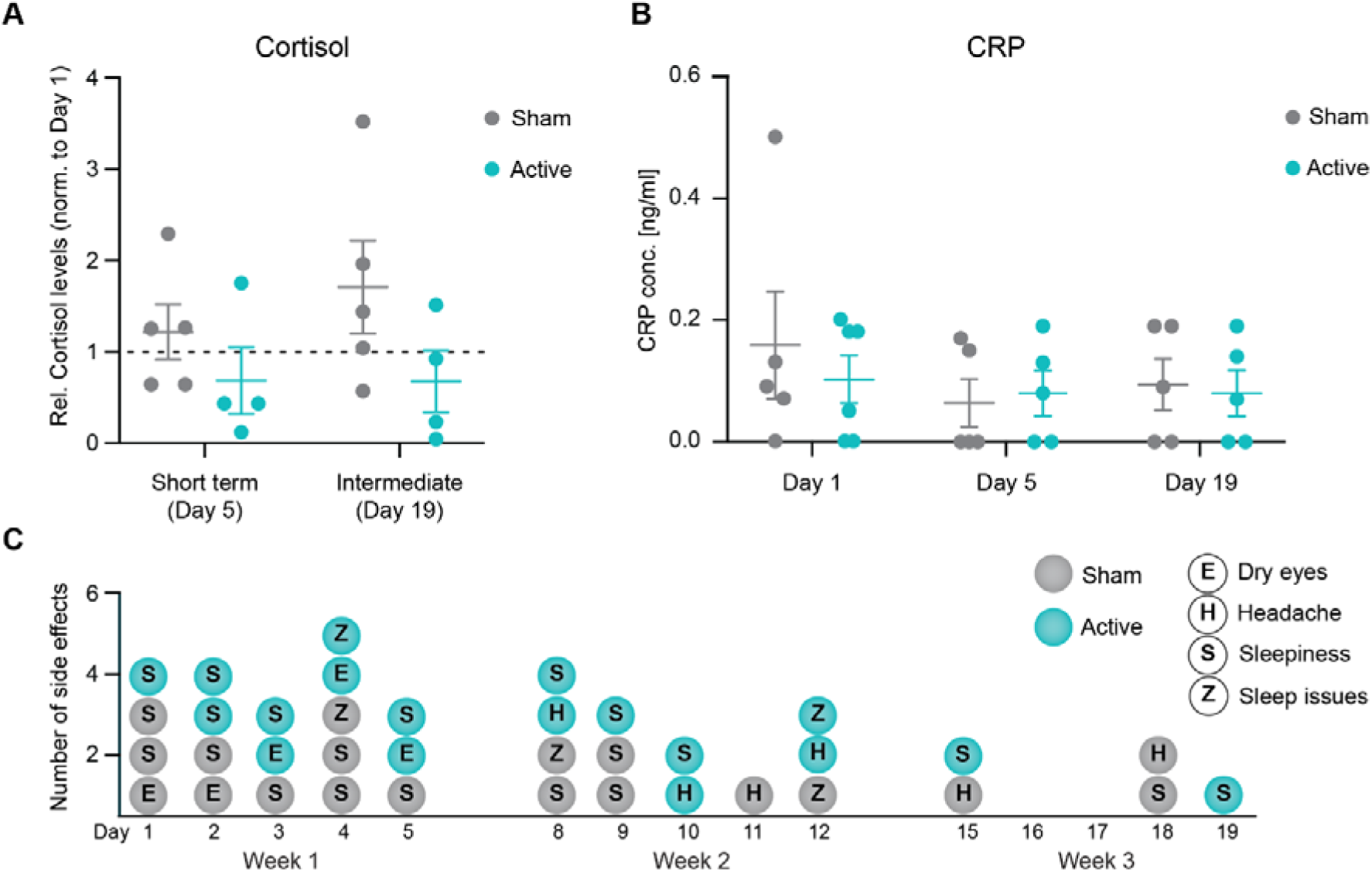
Systemic effects of 60 Hz entrainment. **(A-B)** Molecular marker analysis for stress and inflammatory responses in saliva following 60 Hz light stimulation. **(A)** Competitive ELISA quantification of salivary cortisol levels, normalized to baseline (day 1). Cortisol levels were measured after 1 week (day 5) of stimulation and after 3 weeks of intermediate stimulation (day 19) in the active (turquoise) and sham (gray) group (active: n = 4; sham: n = 5). Data are presented as scatter dot plots showing mean ± standard deviation (SD). Statistical significance was determined using two-way ANOVA with multiple comparisons, followed by the Shapiro-Wilk test. **(B)** Quantification of CRP levels in saliva, measured on day 1, day 5, and day 19 of the study. Participants received either active (turquoise) or sham (gray) stimulation (active: n = 5; sham: n = 5). Data are presented as scatter dot plots showing mean ± standard deviation (SD). Statistical significance was determined using Kruskal-Wallis test multiple comparisons. **C)** day by day record of side effects during the stimulation. Each dot represents a single side effect. No statistically significant differences were observed across groups (see **Table 2**).

### 4.5 Effects on CRP levels measured in saliva

In addition to the neuroendocrine axis, we wondered whether 60 Hz could have other somatic effects. We were particularly interested in potential immune responses, given the body of evidence indicating that both 40 Hz^15^ and 60 Hz^19^ can affect microglial phenotype in preclinical models. As a general marker of immune activation we measured C-reactive protein (CRP) in saliva samples of participants. No significant differences were observed in the levels of CRP on day 1, 5 and 19, when comparing active and sham groups (**Fig 3B**). These findings suggest that 60 Hz stimulation, even over a three-week period, does not induce a detectable inflammatory response.

### 4.6 Tolerability

Neural entrainment using white light is generally safe and well tolerated by subjects with only minor side effects such as headaches and eye strain^17^. Our study was designed to minimize the discomfort experienced by subjects, by using an LED-generated light whose spectrum largely overlaps with daylight (**Supplementary** Fig. 1 **A**), and at a very low intensity which subjects could adjust according to their preference (between 74 and 94 μW -measured at 2cm distance from the LED strip). However, the tolerability of 60 Hz neural entrainment over extended periods has not been thoroughly studied. This is particularly relevant because our protocol induced an artificial and sustained 60 Hz activity pattern over three weeks which could potentially lead to discomfort.

To address this, we assessed the overall tolerability of the procedure through questionnaires, participant self-reports of side effects, and open-ended feedback. Questionnaire data, as well as side effects occurrence and dropouts, were compared statistically between the active and sham groups. The results of this analysis are shown in **Table 2** and **Fig 3C**.

Out of the 14 participants in the study, none found the stimulation, whether sham or active, unpleasant. The stimulation was rated as highly tolerable with an average score of 4.5 in the sham group and 4.8 in the active group. All participants indicated a willingness to undergo the procedure again, with an average score of 3.5 in the sham and 4.0 in the active group. Side effects were recorded in both groups at similar rates and were considered minor.

Consistent with findings from other neural entrainment studies using 40Hz^17,18,32^, the most common side effect was sleepiness, reported by 66.7% of the sham group and 62.5% of the active group, followed by dry eyes/eye strain. Other reported side effects were new onset headache, sleep disturbances; quantification is presented in Table 2.

Interestingly, the number of reported side effects declined over time in both groups (total side effects in week one: 9 in active and 11 in sham group; total side effects in week 2: 7 in active and 6 in sham group; total side effects in week 3: 2 in active and 3 in sham group; see **Figure 3 C**) corresponding to a decrease of 78% and 73 % in active and sham groups, respectively, by week 3 as compared to week 1. Overall, the stimulation appeared to be well tolerated and safe, with no unexpected adverse events recorded.

## 5. Discussion

### 5.1 Technical considerations

While several studies have investigated the effect of 40 Hz neural entrainment, 60 Hz entrainment is relatively less explored. Given the role of high gamma oscillations in cognitive functions^33^, their alterations in several neurological and psychiatric diseases^14^, and the preclinical effects of 60 Hz on neuroplasticity^19^, 60 Hz neural entrainment is a topic of high interest. One study^34^ showed successful and widespread entrainment with acute 60 Hz visual stimulation, but the effect of prolonged stimulation has not been tested. A few studies have used transcranial alternating current stimulation (tACS) at 60 Hz in healthy volunteers and showed entrainment^35^, but the effect of visual stimulation has not been tested. The effects on somatic response of subjects and well-being have not been systematically recorded in most studies.

The scarcity of studies on 60 Hz entrainment might be due to inherent technical challenges in working with this specific frequency. For example in North America, a 60 Hz EEG signal overlaps with electrical noise (voltage of 110 V and a frequency of 60 Hz), making it impossible to analyze the 60 Hz component and frequencies close to it. Being based in Europe, we worked with a voltage of 220 V at 50Hz, therefore the electrical noise could be easily filtered out and did not interfere with our analysis. Another technical aspect that makes this type of work challenging is the interference between the electrical noise of the flickering LEDs and the EEG recording. To overcome this issue, we have encased the LED stripe in a mini Faraday cage. This resulted in a clear reduction of electrical noise as measured with multifield-EMF. The fact that the 60 Hz entrainment signal in EEG declines over time in each subject, while the LED light was always active throughout the 3 weeks, demonstrates that we were successful in removing noise and that the observed signal is driven by brain activity.

### 5.2 60 Hz neural entrainment power

To our knowledge, this is the first study to combine detailed EEG signal analysis over a three-week period with assessments of somatic and quality-of-life responses in healthy volunteers receiving light-induced neural entrainment.

The presence of the distinct 60 Hz peak in the PSD of active group participants confirmed that our intermittent light stimulation effectively entrained neural activity at the stimulation frequency. This finding aligns with previous studies demonstrating frequency-specific entrainment using acute stimulation with visual stimuli^37^, as well as rhythmic sensory stimulation, particularly at gamma frequencies such as 40 Hz^11,38^.

Further EEG analysis confirmed a strong entrainment in the active group, which was not limited to a few individuals or specific channels. The normalized power of individual channels ranged from 0.8 to 12.9 on day 1, 0.7 to 9.3 on day 5, and 0.7 to 3.6 on day 19. As shown in **Fig. 1F**, most channels exhibited entrainment. Subject-specific responses, presented in **Fig. 1F and Supplementary** Fig. 1H, revealed considerable variability across individuals; however, the overall response was consistent across subjects. The STFT analysis (**Fig. 1G**) further demonstrated that entrainment was sustained over the 20-minute stimulation period for each individual.

### 5.3 60 Hz neural entrainment topographic distribution

Next, we investigated the topographical distribution of the entrainment. Interestingly, we did not observe a focal neural activation limited to the visual cortex (the primary target of visual stimulation) but rather an activity that spread to the rest of the cortex, including the parietal cortex, reaching the frontal cortex. Similar spreading has been observed with other entrainment modalities such as 40 Hz light^32^ and transcranial electrical stimulation^39^. Such a diffuse pattern of brain activity could be driven by synaptic connections or traveling cortical waves. Indeed, gamma entrainment has been shown to induce traveling waves connecting the occipital/parietal regions to the frontal region of the cortex through the temporal region^40^.

In addition, we found that the brain activation pattern is not random but is highly synchronized, as revealed by our PLV analysis. This finding is consistent with reports that flickering simulations increase phase connectivity between brain regions^41^. While the mechanism underlying such connectivity effect is not known, it is likely to involve functional neuroplastic changes.

### 5.4 Effect of time

In line with the idea of neuroplastic changes occurring with 60 Hz stimulation, we observed a neural adaptation to the entrainment over time, namely after 5 and 15 days of stimulation (corresponding to day 5 and day 19).

Our results indicate that the response is highest on day 1 during the very first stimulation, still present at day 5 but lower, and further reduced at day 19. For instance, the average normalized power across all channels of all subjects went from 2.8 on day 1 (SD 2.69) to 2.4 on day 5 (SD 1.94) and 1.54 on day 19 (SD 0.61). All features of the 60 Hz entrainment that we analysed (the normalized entrainment power (**Fig 1F**), the topographic distribution (**Fig 1E**) and the PLV analysis (**Fig 2A,B**) showed a similar pattern.

The reduction in entrainment power as well as PLV over time suggests a neural adaptation to the repeated stimulations. Similar adaptation has been previously described with repeated acoustic stimulations^42^ as well as with 40 Hz combined visual and auditory entrainment^32^. The adaptation is likely related to homeostatic plasticity^43^. Briefly, repeated stimulations will first stimulate Hebbian plasticity and the formation of new connections (quick response in minutes/hours); this is followed, at a slower timescale (days) by homeostatic plasticity which removes connections to assure stability^44^. It is intriguing to observe that, in animal experiments, 60 Hz was shown by our group to promote juvenile-like neuroplasticity via remodeling of the PNN^19^. PNN remodeling might be the molecular underpinning of the functional synaptic changes observed in healthy volunteers receiving 60 Hz flickering light; this hypothesis remains to be investigated.

### 5.5 Somatic response and tolerability

To the best of our knowledge this is the first study to characterize the somatic response to 60 Hz flickering light. Evidence indicates that other NIBS can modulate the neuroendocrine axis^29,30^ and can have an impact on microglia, the resident immune cells of the brain^15,19^. We were therefore interested in studying not just the brain response to 60 Hz flickering light, but also the somatic response. Importantly, we show that 60 Hz in healthy humans does not elicit a strong immune response, as CRP levels appear not statistically different across groups (**Fig 3B**). Similarly, the strong brain activity response we observed was not accompanied by any sign of stress, as measured by cortisol levels (**Fig 3A**); if anything, we observed a trend towards reduction. Considering the small sample size, the effect of 60 Hz on cortisol might deserve further investigation in a larger experiment.

Confirming the indication coming from the cortisol measurements, 60 Hz stimulation was overall well tolerated by the participants (**Fig 3C**). Based on the literature (mostly on 40 Hz stimulations^17,18,32^), we expected only minor side effects. Indeed, the vast majority of participants reported some side effects (listed in **Table 2**; eye strain, sleepiness, new onset headache and sleep disturbances), but all of them, in both sham and active groups, were minor and did not require medical assistance. The responses the participants gave to our questionnaire confirmed that the stimulation (both active and sham) was well tolerated, and all would have done it again; the two dropouts we had were due to personal reasons, and none of them was stimulation-related. Nobody asked to interrupt or shorten a stimulation session. Since this was a particularly young and healthy research cohort, it remains to be assessed whether the 60 Hz effect is the same in a more heterogenous population. However, these results confirm that the EEG changes we observed were not confounded by stress or discomfort caused by the stimulation itself.

### 5.7 Limitations

This pilot study has several limitations. First, sample size likely limited our ability to detect additional differences at group level. Due to insufficient statistical power, we were unable to explore potential influences of sex, gender, age, and time of day on the neural response to 60 Hz flickering light, which remain important areas for future research. Nevertheless, despite the low number of subjects, our findings clearly demonstrate neural entrainment effects and its evolution over time.

Second, while we did our best to ensure a balance for sex and age, the participants of this study were all very young, highly educated and mostly white. The neural entrainment following 60 Hz flickering light in a more diverse population remains to be established.

Third, the use of 8-channel EEG, gave us very low spatial resolution and did not allow us to study in detail the topographical distribution of the entrainment.

## 6. Conclusions

Externally-induced neural entrainment with 60 Hz flickering light over 3 weeks is strong, synchronized across brain regions, and leads to neural habituation, with no major side effects. Several FDA-approved drugs for psychiatric conditions, including SSRIs and ketamine, boost neuroplasticity^45^, promote the remodeling of extracellular matrix^46^, and induce gamma activity^47^. Externally induced 60 Hz entrainment might therefore be a new approach for modulating brain activity and inducing neuroplasticity, with implications for our basic understanding of brain physiology as well as treatment of psychiatric disorders. Since light is a non-invasive, easily implementable, amenable to at-home treatment, and relatively cheap methodology, this research area warrants further investigation.

## Data availability

Data is available upon request to the corresponding author.

## 7 Acknowledgments and funding sources

This study was funded by Syntropic Medical and supported by an Austria Wirtschaftsservice (AWS) grant (grant number P2414247 to Syntropic Medical).

The authors thank all the participants for their time and commitment to this study. Special thanks to Dr. Verena Seiboth, Mrs. Verena Schmied, and Dr. Katalin Szigeti for their valuable assistance.

## 9 CRediT authorship contribution statement

MohammadAmin Alamalhoda (conceptualization, methodology, software, validation, formal analysis, investigation, data curation, writing original draft, visualization), Friederike Leesch (methodology, validation, formal analysis, investigation, data curation, writing original draft, visualization), Francesca Giovanetti (investigation, review and editing, project administration), Eoghan Dunne (review and editing), Giuseppina Pilloni (review and editing), Mark Caffrey (conceptualization, resources, funding acquisition, review and editing) Jack O’Keeffe (conceptualization, resources, funding acquisition, review and editing), Alessandro Venturino (conceptualization, methodology, validation, investigation, writing original draft, review and editing, supervision, funding acquisition), Maria Teresa Ferretti (conceptualization, methodology, investigation, formal analysis, writing original draft, review and editing, supervision).

---

**Supplementary Figure 1.**
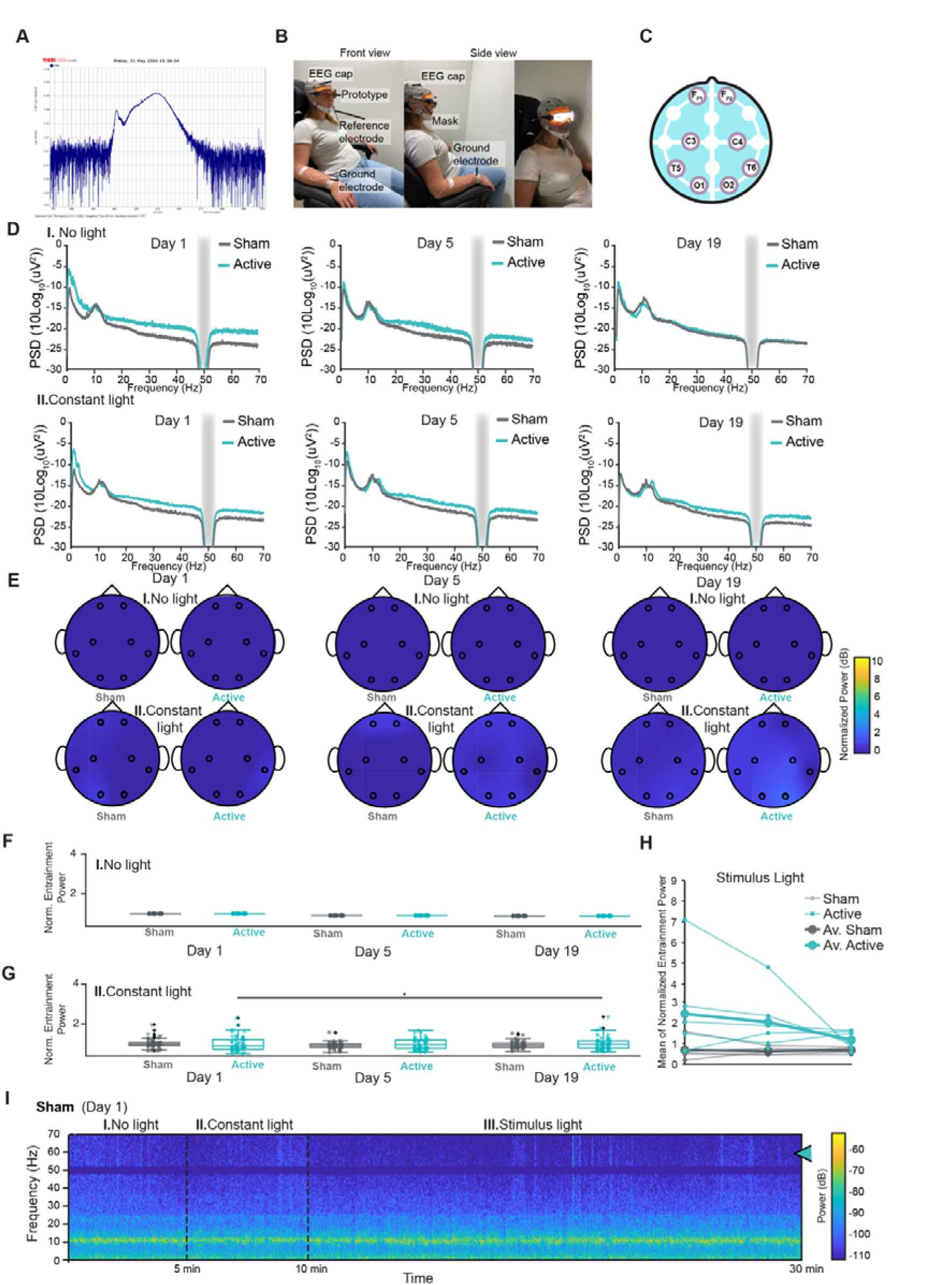
Experimental set up and control conditions to assess 60 Hz-induced brain entrainment. **(A**) Graph showing the light spectrum of the LEDs ranging from 440 nm to 770 nm, similar to daylight wavelengths. **(B**) Experimental setup on EEG days: subjects were seated on a chair during the stimulation (picture of one of the authors). (**C**) The figure illustrates the location of the 8 EEG channels utilized in the study, mapped according to the 10-20 electrode placement system. **(D**) Scalp EEG power spectral density (PSD) averaged across all channels for participants in each group under (I.) No light and (II.) Constant light conditions . The gray bar indicates the 50 Hz line noise, which was notch-filtered. (**E**) Topographic maps showing normalized changes in 60 Hz PSD (relative to baseline) averaged across participants of each group under (I.) No light and (II.) Constant light conditions. (**F, G**) Significant differences in normalized changes in 60 Hz PSD values were observed across all channels between the active and sham groups under (I.) No light and (II.) Constant light conditions on days 1, 5, and 19. Statistical significance for these inter-group comparisons was assessed using the Wilcoxon rank-sum test. Furthermore, within the active group, significant differences in normalized PSD were detected between day 1 vs. day 5 and day 1 vs. day 19, accounting for repeated measurements. These intra-group comparisons across days were evaluated using the Kruskal-Wallis test, followed by post-hoc pairwise comparisons performed using Dunn’s test with Bonferroni correction. The normality of the data was assessed using the Shapiro-Wilk test, and non-parametric methods were applied due to deviations from normality. In this figure, different channels are shape-coded. Only significant differences are indicated, with * indicating p < 0.05 and *** indicating p < 0.001. Detailed p-values are provided in **Supplementary Table 2**. (**H**) The figure displays the average normalized PSD of all channels for each subject in both groups over days 1, 5, and 19, represented as individual lines. The thick line indicates the group average across all channels and subjects. Data were recorded during light stimulation, with 60 Hz flickering light for the active group and constant light for the sham group.**(I**) Short-Time Fourier Transform (STFT) of a representative sham group participant, demonstrating no visible 60 Hz indicated by the turquoise arrow.

**Supplementary Figure 2.**
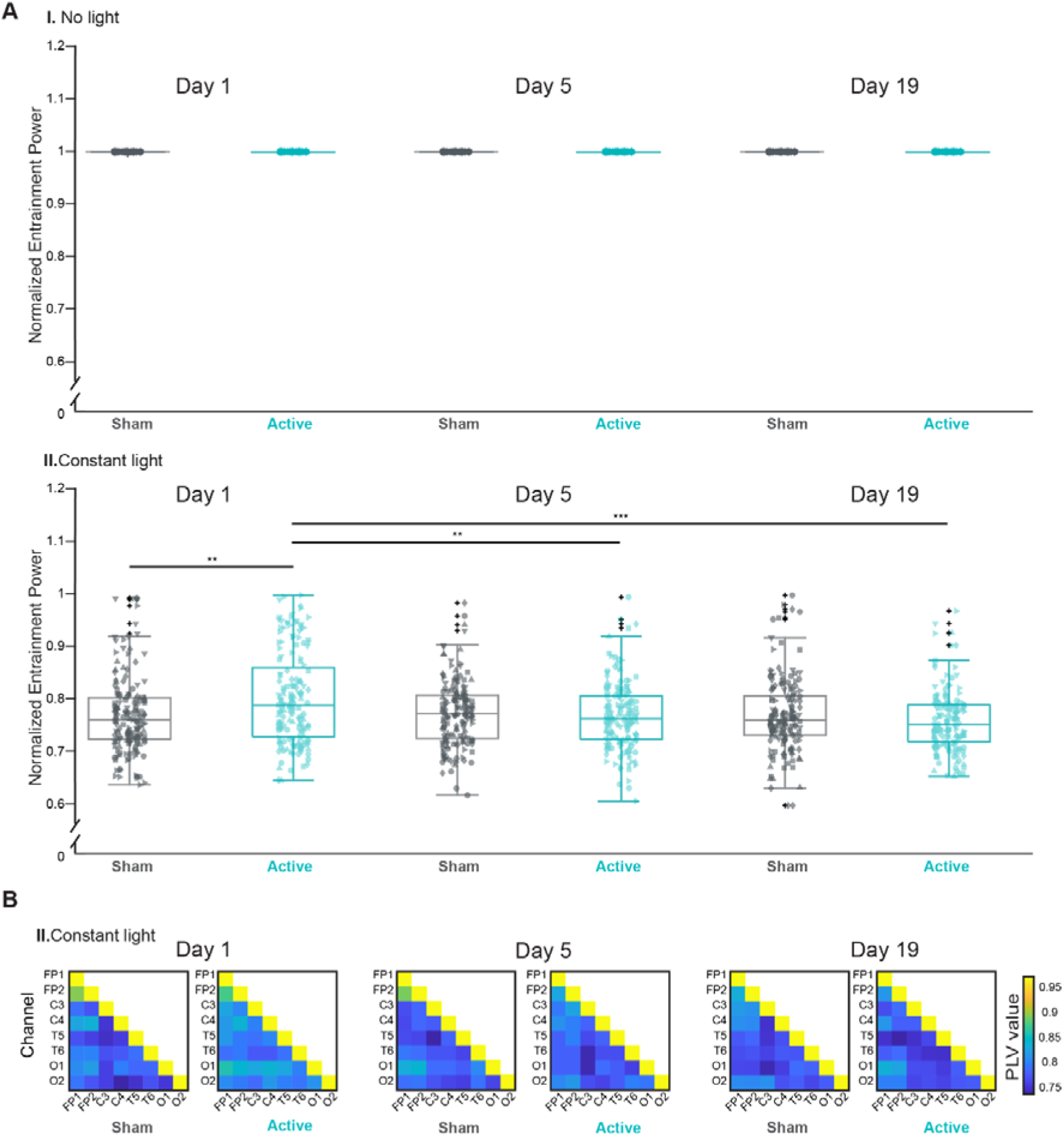
Assessment of synchronisation of brain activity during 60 Hz entrainment during no light and constant light stimulus. (**A**) No significant differences in normalized PLV (normalized to PLV matrix of No Light condition) were observed across all channel pairs between the active and sham groups under (I.) No light and (II.) Constant light conditions on days 1, 5, and 19. Inter-group comparisons were assessed using the Wilcoxon rank-sum test. Intra-group comparisons across days were evaluated using the Kruskal-Wallis test, followed by post-hoc pairwise comparisons performed using Dunn’s test with Bonferroni correction. Normality of the data was assessed using the Shapiro-Wilk test, and non-parametric methods were employed due to deviations from normality. (**B**) PLV matrices for the active and sham groups across experimental days under (I.) No light and (II.) Constant light conditions. Each element in the matrix represents the PLV for specific pairs of EEG channels, with diagonal elements showing a value of 1, indicating PLV between identical signals. Significant differences are indicated, with ** indicating p < 0.01 and *** indicating p < 0.001. Detailed p-values are provided in **Supplementary Table 2.**

## References

1. Davidson, B. et al. Neuromodulation techniques – From non-invasive brain stimulation to deep brain stimulation. Neurotherapeutics 21, e00330 (2024).

2. Fitzsimmons, S. M. D. D., Oostra, E., Postma, T. S., Van Der Werf, Y. D. & Van Den Heuvel, O. A. Repetitive Transcranial Magnetic Stimulation–Induced Neuroplasticity and the Treatment of Psychiatric Disorders: State of the Evidence and Future Opportunities. Biol. Psychiatry 95, 592– 600 (2024).

3. Pirnia, T. et al. Electroconvulsive therapy and structural neuroplasticity in neocortical, limbic and paralimbic cortex. Transl. Psychiatry 6, e832 (2016).

4. Kunze, T., Hunold, A., Haueisen, J., Jirsa, V. & Spiegler, A. Transcranial direct current stimulation changes resting state functional connectivity: A large-scale brain network modeling study. NeuroImage 140, 174–187 (2016).

5. Pan, F. et al. Effects of neuronavigation-guided rTMS on serum BDNF, TrkB and VGF levels in depressive patients with suicidal ideation. J. Affect. Disord. 323, 617–623 (2023).

6. Cohen, S. L., Bikson, M., Badran, B. W. & George, M. S. A visual and narrative timeline of US FDA milestones for Transcranial Magnetic Stimulation (TMS) devices. Brain Stimulat. 15, 73–75 (2022).

7. Voelker, R. Brain Stimulation Approved for Obsessive-Compulsive Disorder. JAMA 320, 1098 (2018).

8. Yaakub, S. N. et al. Transcranial focused ultrasound-mediated neurochemical and functional connectivity changes in deep cortical regions in humans. Nat. Commun. 14, 5318 (2023).

9. Violante, I. R. et al. Non-invasive temporal interference electrical stimulation of the human hippocampus. Nat. Neurosci. 26, 1994–2004 (2023).

10. Li, Z. et al. Transcranial low-level laser stimulation in the near-infrared-II region (1064 nm) for brain safety in healthy humans. Brain Stimulat. 17, 1307–1316 (2024).

11. Thut, G., Schyns, P. G. & Gross, J. Entrainment of Perceptually Relevant Brain Oscillations by Non-Invasive Rhythmic Stimulation of the Human Brain. Front. Psychol. 2, (2011).

12. Kahana, M. J. The Cognitive Correlates of Human Brain Oscillations. J. Neurosci. 26, 1669– 1672 (2006).

13. Tada, M. et al. Alterations of auditory-evoked gamma oscillations are more pronounced than alterations of spontaneous power of gamma oscillation in early stages of schizophrenia. Transl. Psychiatry 13, 218 (2023).

14. Guan, A. et al. The role of gamma oscillations in central nervous system diseases: Mechanism and treatment. Front. Cell. Neurosci. 16, 962957 (2022).

15. Iaccarino, H. F. et al. Gamma frequency entrainment attenuates amyloid load and modifies microglia. Nature 540, 230–235 (2016).

16. Martorell, A. J. et al. Multi-sensory Gamma Stimulation Ameliorates Alzheimer’s-Associated Pathology and Improves Cognition. Cell 177, 256–271.e22 (2019).

17. Hajós, M. et al. Safety, tolerability, and efficacy estimate of evoked gamma oscillation in mild to moderate Alzheimer’s disease. Front. Neurol. 15, 1343588 (2024).

18. He, Q. et al. A feasibility trial of gamma sensory flicker for patients with prodromal Alzheimer’s disease. Alzheimers Dement. Transl. Res. Clin. Interv. 7, e12178 (2021).

19. Venturino, A. et al. Microglia enable mature perineuronal nets disassembly upon anesthetic ketamine exposure or 60-Hz light entrainment in the healthy brain. Cell Rep. 36, 109313 (2021).

20. Report of the committee on methods of clinical examination in electroencephalography. Electroencephalogr. Clin. Neurophysiol. 10, 370–375 (1958).

21. Delorme, A. & Makeig, S. EEGLAB: an open source toolbox for analysis of single-trial EEG dynamics including independent component analysis. J. Neurosci. Methods 134, 9–21 (2004).

22. Jung, T. P. et al. Removing electroencephalographic artifacts by blind source separation. Psychophysiology 37, 163–178 (2000).

23. Oppenheim, A. & Shafer, R. Discrete-Time Signal Processing. (Prentice Hall, 1999).

24. cohen, l. Time-Frequency Analysis. (Prentice Hall, 1995).

25. Lachaux, J.-P., Rodriguez, E., Martinerie, J. & Varela, F. J. Measuring phase synchrony in brain signals. Hum. Brain Mapp. 8, 194–208 (1999).

26. de Almeida, P. D. V., Grégio, A. M. T., Machado, M. A. N., de Lima, A. A. S. & Azevedo, L. R. Saliva composition and functions: a comprehensive review. J. Contemp. Dent. Pract. 9, 72–80 (2008).

27. Reber, T. P. et al. Single-neuron mechanisms of neural adaptation in the human temporal lobe. Nat. Commun. 14, 2496 (2023).

28. Homann, J., Koay, S. A., Chen, K. S., Tank, D. W. & Berry, M. J. Novel stimuli evoke excess activity in the mouse primary visual cortex. Proc. Natl. Acad. Sci. 119, e2108882119 (2022).

29. Evers, S., Hengst, K. & Pecuch, P. W. The impact of repetitive transcranial magnetic stimulation on pituitary hormone levels and cortisol in healthy subjects. J. Affect. Disord. 66, 83–88 (2001).

30. Brunoni, A. R. et al. Polarity- and valence-dependent effects of prefrontal transcranial direct current stimulation on heart rate variability and salivary cortisol. Psychoneuroendocrinology 38, 58–66 (2013).

31. Menshov, V. A. et al. Influence of Nicotine from Diverse Delivery Tools on the Autonomic Nervous and Hormonal Systems. Biomedicines 10, 121 (2022).

32. Chan, D. et al. Gamma frequency sensory stimulation in mild probable Alzheimer’s dementia patients: Results of feasibility and pilot studies. PLOS ONE 17, e0278412 (2022).

33. Griffiths, B. J. & Jensen, O. Gamma oscillations and episodic memory. Trends Neurosci. 46, 832–846 (2023).

34. Jones, M. et al. Gamma Band Light Stimulation in Human Case Studies: Groundwork for Potential Alzheimer’s Disease Treatment. J. Alzheimers Dis. 70, 171–185 (2019).

35. Hanley, C. J., Singh, K. D. & McGonigle, D. J. Transcranial modulation of brain oscillatory responses: A concurrent tDCS–MEG investigation. NeuroImage 140, 20–32 (2016).

36. Notbohm, A., Kurths, J. & Herrmann, C. S. Modification of Brain Oscillations via Rhythmic Light Stimulation Provides Evidence for Entrainment but Not for Superposition of Event-Related Responses. Front. Hum. Neurosci. 10, (2016).

37. Khachatryan, E. et al. Cognitive tasks propagate the neural entrainment in response to a visual 40 Hz stimulation in humans. Front. Aging Neurosci. 14, 1010765 (2022).

38. Herrmann, C. S., Munk, M. H. J. & Engel, A. K. Cognitive functions of gamma-band activity: memory match and utilization. Trends Cogn. Sci. 8, 347–355 (2004).

39. Kar, K. & Krekelberg, B. Transcranial electrical stimulation over visual cortex evokes phosphenes with a retinal origin. J. Neurophysiol. 108, 2173–2178 (2012).

40. Alamalhoda, M. A., Lahijanian, M., Aghajan, H. & Vahabi, Z. Traveling waves induced by gamma entrainment can explain its therapeutic effects for Alzheimer’s disease. Alzheimers Dement. 19, e066261 (2023).

41. Lahijanian, M., Aghajan, H. & Vahabi, Z. Auditory gamma-band entrainment enhances default mode network connectivity in dementia patients. Sci. Rep. 14, 13153 (2024).

42. Todorovic, A., Van Ede, F., Maris, E. & De Lange, F. P. Prior Expectation Mediates Neural Adaptation to Repeated Sounds in the Auditory Cortex: An MEG Study. J. Neurosci. 31, 9118– 9123 (2011).

43. Menicucci, D., Lunghi, C., Zaccaro, A., Morrone, M. C. & Gemignani, A. Mutual interaction between visual homeostatic plasticity and sleep in adult humans. eLife 11, e70633 (2022).

44. Fauth, M. & Tetzlaff, C. Opposing Effects of Neuronal Activity on Structural Plasticity. Front. Neuroanat. 10, (2016).

45. Klöbl, M. et al. Escitalopram modulates learning content-specific neuroplasticity of functional brain networks. NeuroImage 247, 118829 (2022).

46. Diniz, C. R. A. F. et al. Fluoxetine and Ketamine Enhance Extinction Memory and Brain Plasticity by Triggering the p75 Neurotrophin Receptor Proteolytic Pathway. Biol. Psychiatry S0006322324014252 (2024) doi:10.1016/j.biopsych.2024.06.021.

47. Adam, E., et al. Ketamine can produce oscillatory dynamics by engaging mechanisms dependent on the kinetics of NMDA receptors. Proc. Natl. Acad. Sci. 121, e2402732121 (2024).

