## Supplementary Table 1 for "Flickering white light stimulation at 60 Hz induces strong, widespread neural entrainment and synchrony in healthy subjects"

| **Inclusion Criteria** | - Participant is male or female or diverse, between the ages of 18 – 65 - Participant is willing and able to sign informed consent form - Participant has signed informed consent form |
| --- | --- |
| **Exclusion Criteria** | - Participants who are pregnant. - Diagnosis of neurodegenerative diseases or psychiatric or brain injury/stroke, in particular:   - -history of seizure or epilepsy   - -known diagnosis of migraine   - -tinnitus - Active treatment with Clopidrogel - Photosensitivity - Retinal diseases or cataract - Autoimmune diseases - Any skin lesion or skull morphology that might interfere with proper EEG reading - Participants with a work relationship with the Siegert lab or any of the sponsor cofounders |

**Supplemntary Table 1 (S1)**

**Inclusion and exclusion criteria**
