## Supplementary figures and images for "Flickering white light stimulation at 60 Hz induces strong, widespread neural entrainment and synchrony in healthy subjects"

### Supplementary Figure 1

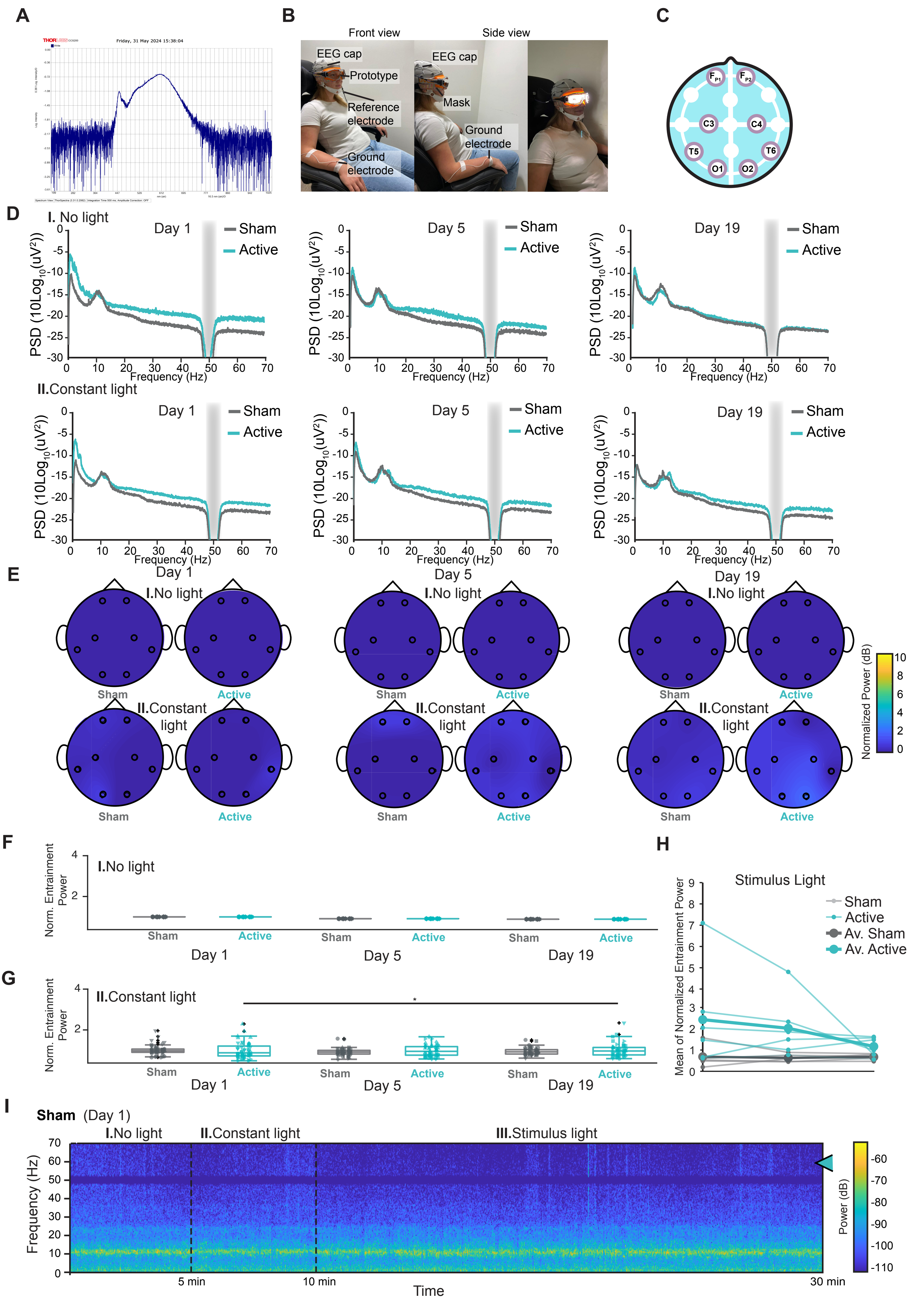

### Supplementary Figure 2

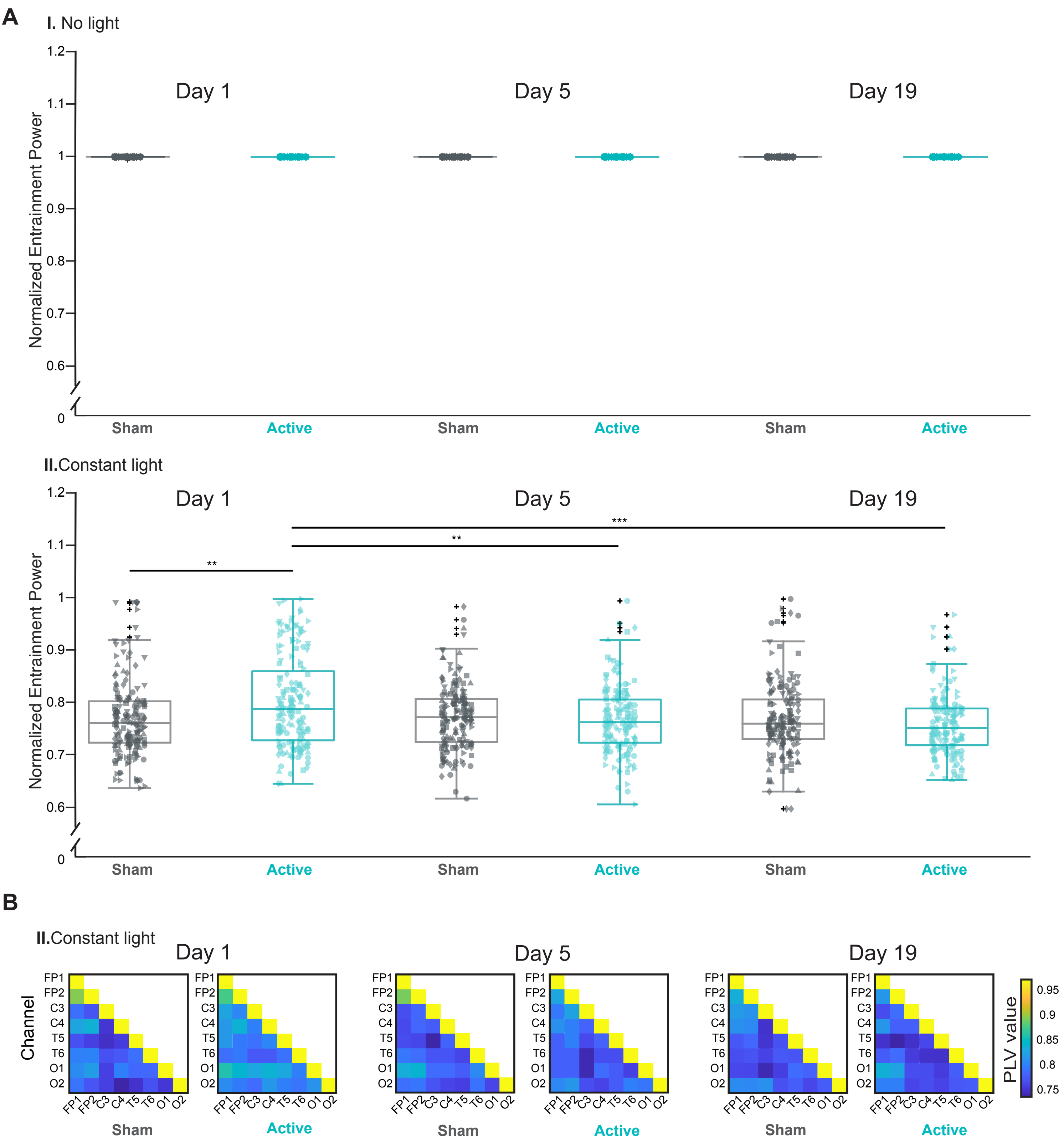
